# Determinants of genetic structure of the Sub-Saharan parasitic wasp *Cotesia sesamiae*

**DOI:** 10.1101/194084

**Authors:** Antoine Branca, Bruno Le Ru, Paul-André Calatayud, Julius Obonyo, Boaz Muzyoka, Claire Capdevielle-Dulac, Laure Kaiser-Arnauld, Jean-François Silvain, Jérémy Gauthier, Corentin Paillusson, Philippe Gayral, Elisabeth A. Herniou, Stéphane Dupas

## Abstract

Parasitoid life style represents one of the most diversified life history strategies on earth. There are however very few studies on the variables associated with intraspecific diversity of parasitoid insects, especially regarding the relationship with spatial, biotic and abiotic ecological factors. *Cotesia sesamiae* is a Sub-Saharan stenophagous parasitic wasp that parasitizes several African stemborer species with variable developmental success. The different host-specialized populations are infected with different strains of *Wolbachia*, an endosymbiotic bacterium widespread in arthropods that is known for impacting life history traits notably reproduction, and consequently species distribution. In this study, first we analyzed the genetic structure of *C. sesamiae* across Sub-Saharan Africa, using 8 microsatellite markers, and 3 clustering software. We identified five major population clusters across Sub-Saharan Africa, which probably originated in East African Rift region and expanded throughout Africa in relation to host genus and abiotic factors such as climatic classifications. Using laboratory lines, we estimated the incompatibility between the different strains of *Wolbachia* infecting *C. sesamiae*. We observed an incompatibility between *Wolbachia* strains was asymmetric; expressed in one direction only. Based on these results, we assessed the relationships between direction of gene flow and *Wolbachia* infections in the genetic clusters. We found that *Wolbachia-*induced reproductive incompatibility was less influential than host specialization in the genetic structure. Both *Wolbachia* and host were more influential than geography and current climatic conditions. These results are discussed in the context of African biogeography, and co-evolution between *Wolbachia*, virus parasitoid and host, in the perspective of improving biological control efficiency through a better knowledge of the biodiversity of biological control agents.

## Introduction

Understanding the extraordinary biodiversity of insects requires both analyzing large scale beta diversity patterns (Heino *et al.* 2015) and unraveling mechanisms of genetic differentiation among populations including geographic, abiotic or biotic interactions (Roderick 1996). Parasitoid wasps are one of the most diverse groups of insects (Grimaldi 2005). Coevolutionary interactions are likely major diversifying forces in host–parasitoid systems due to the strength of reciprocal selection pressures (Van Valen 1973; Henry *et al.* 2008). As strong insect antagonists, they are the most used agents for biological control programs, which provide one of the best alternatives to chemical control of insect pests (Harvey 2011). There are theoretical expectations that host parasitoid coevolution generates diversity because several traits related to host specificity, such as specific virulence and host recognition, are mechanistically linked to reproductive isolation, especially when the parasitoid mates on the host just after emergence (Dupas *et al.* 2008; Hoskin & Higgie 2010). Other biotic interactions, particularly those involving microorganisms affecting reproduction such as *Wolbachia* sp., are expected to drive diversification of parasitoids (Bordenstein *et al.* 2001; Branca *et al.* 2009). To distinguish between the different ecological factors responsible for population structure, a combination of, on the one hand, laboratory data on reproductive incompatibility and, on the other hand, field data on the geographic structure of ecological drivers and population differentiation are needed.

### Cotesia sesamiae

Cameron (Hymenoptera: Braconidae) is a parasitoid wasp widespread in Sub-Saharan Africa that has been used in biological control for controlling *Busseola fusca* (Fuller) (Lepidoptera: Noctuidae), a major stemborer pest of maize and sorghum crops (Kfir 1995; Kfir *et al.* 2002). *Cotesia sesamiae* is a stenophagous parasitoid that successfully parasitizes diverse host species (Ngi-Song *et al.* 1995; Branca *et al.* 2011). However, a variation in parasitism success on different hosts has been shown among populations of parasitoids (Mochiah *et al.* 2002a; Gitau *et al.* 2010). In contrast to the *C. sesamiae* population from Mombasa - coastal Kenya (avirulent towards *B. fusca*), the *C. sesamiae* population from Kitale – inland Kenya (virulent towards *B. fusca*) is able to develop in *B. fusca*, but both develop in *Sesamia calamistis* Hampson (Lepidoptera: Noctuidae), the main host of *C. sesamiae* in coastal Kenya (Ngi-Song *et al*. 1995). These differences in host acceptance and development have been linked to the observed polymorphism of a candidate gene, CrV1, located on the bracovirus locus (Dupas *et al.* 2008; Gitau *et al.* 2007; Branca *et al.* 2011). Bracoviruses are symbiotic polydnaviruses integrated to the genome of braconid wasps, contributing to their adaptive radiations (Whitfield 2002; Dupuy *et al.* 2006). The viruses constitute the major components of the calyx fluid of the wasp and are expressed in parasitoid host cells, regulating its physiology, development and immunology (Beckage 1998). In particular, the CrV1 gene, has been shown to contribute to immune suppression by active de-structuration of the cytoskeleton of host immune cells (Asgari *et al.* 1997). A comparative genomics study of the virus between *Cotesia* species and *C. sesamiae* populations, virulent and avirulent against *B. fusca*, showed patterns suggesting important role for positive selection, gene duplication and recombination among viral genes in the adaptive diversification process (Jancek *et al.* 2013). Whilst host resistance puts likely a strong selective pressure on local adaptation of the wasp, other ecological and geographic factors must be considered and analyzed for the development of scenario of *C. sesamiae* response to environmental changes. Climatic differences or geographical barriers might weaken the capacity of some *C. sesamiae* populations to colonize areas where the most prevalent host is suitable for parasitoid larval development, even if parasitic wasps have been shown to disperse quite efficiently, sometimes beyond the capacity of their associated host (Antolin & Strong 1987; Ode *et al.* 1998; Van Nouhuys & Hanski 2002; Assefa *et al*. 2008, Santos & Quicke 2011). Other factors such as *Wolbachia* might act as a barrier to gene flow through reproductive incompatibility (Werren 1997; Jaenike *et al.* 2006), which can be especially problematic in the context of biological invasions by preventing crosses between ecological or geographic populations along the range expanded area of the invasive pest host. *Wolbachia* is a widespread bacterium infecting the majority of insect species that can induce reproductive incompatibilities (Werren 1997; Hilgenboecker *et al.* 2008). Several *Wolbachia* strains have been identified in *C. sesamiae* expressing cytoplasmic incompatibilities (CI) between populations of parasitoids (Mochiah *et al.* 2002b). The different populations of *C. sesamiae*, virulent and avirulent against *B. fusca*, are infected with different strains of *Wolbachia* (Branca *et al.* 2011). Reproductive isolation can prevent adapted parasitoid populations to expand across their host range, a phenomenon that could be particularly relevant in biological control programs. In this study, our objective is to analyze the relative importance of neutral geographic factors and major selective forces, biotic (*i.e.* host species and *Wolbachia* strain), abiotic (*i.e.* climate) shaping the genetic structure of the parasitoid *Cotesia sesamiae* across Sub-Saharan Africa. First, the genetic structure was assessed using 8 microsatellites markers with several genetic clustering approaches, each using different pertinent hypotheses in an effort to reach the broadest picture of the structure. Second, we tested the cross incompatibility between differentially *Wolbachia*-infected *C. sesamiae* populations to infer their potential influence on limiting gene flow. Third, we estimated the amount and direction of gene flow in between genetic clusters of selected *C. sesamiae* populations to see if *Wolbachia* infection can affect the parasitoid metapopulation dynamics. Finally, we interpreted geographic patterns of *C. sesamiae* genetic structure in the context of African climate, *Wolbachia* infection and host occurrence.

## Material and methods

### Insects field collection

Infected stemborer larvae were collected in 142 localities of 9 sub-Saharan African countries. GPS positions were recorded at each locality. Stemborer larvae were identified using a larval picture library (corresponding to adult moth identifications), and according to the host plant, as most stemborers are host plant specific (Le Ru *et al.* 2006). Larvae collected from the field were reared on an artificial diet (Onyango & Ochieng’-Odero 1994) until pupation or emergence of parasitoid larvae. After the emergence of cocoons, adult parasitoids were kept in absolute ethanol. Morphological identification of parasitoids was based on genitalia shape following the method of Kimani-Njogu *et al.* (1997). Total genomic DNA of one female per progeny was extracted using the DNeasy Tissue Kit (QIAGEN). If only male were present then analyses were performed on one male. Because wasps are haplodiploids, the haploid genotype of males was converted to homozygous diploids for analyses to avoid discarding data. This should not bias the results because of a very low level of heterozygosity due to very high inbreeding. Total genotyped individuals were 590 females and 47 males discarding individuals with too many missing genotypes (more than 2 over 8 loci).

### Insects rearing

For crossing experiments, females of both virulent and avirulent *C. sesamiae* strains against *B. fusca* were obtained from laboratory-reared colonies. The virulent, thereafter named Kitale (Kit) *C. sesamiae* strain was obtained from *B. fusca* larvae collected from maize fields in Kitale, Western Kenya, in 2006, while the avirulent *C. sesamiae* strain thereafter named Mombasa (Mbsa), was obtained from *S. calamistis* larvae collected from maize fields in coastal Kenya in 2007. The two lines have different *Wolbachia* infection status: Kitale line is infected with *Wolbachia* WCsesB1 strain while Mombasa line is infected with two strains of *Wolbachia* WCsesA and WCsesB2 (Table 1). Twice a year, both colonies were rejuvenated by field collected parasitoids. The wasps of both strains were continuously reared on larvae of *S. calamistis* as previously described Overholt *et al.* (1994). Parasitoid cocoons were kept in Perspex cages (30 x 30 x 30 cm) until emergence.

**Table 1.**
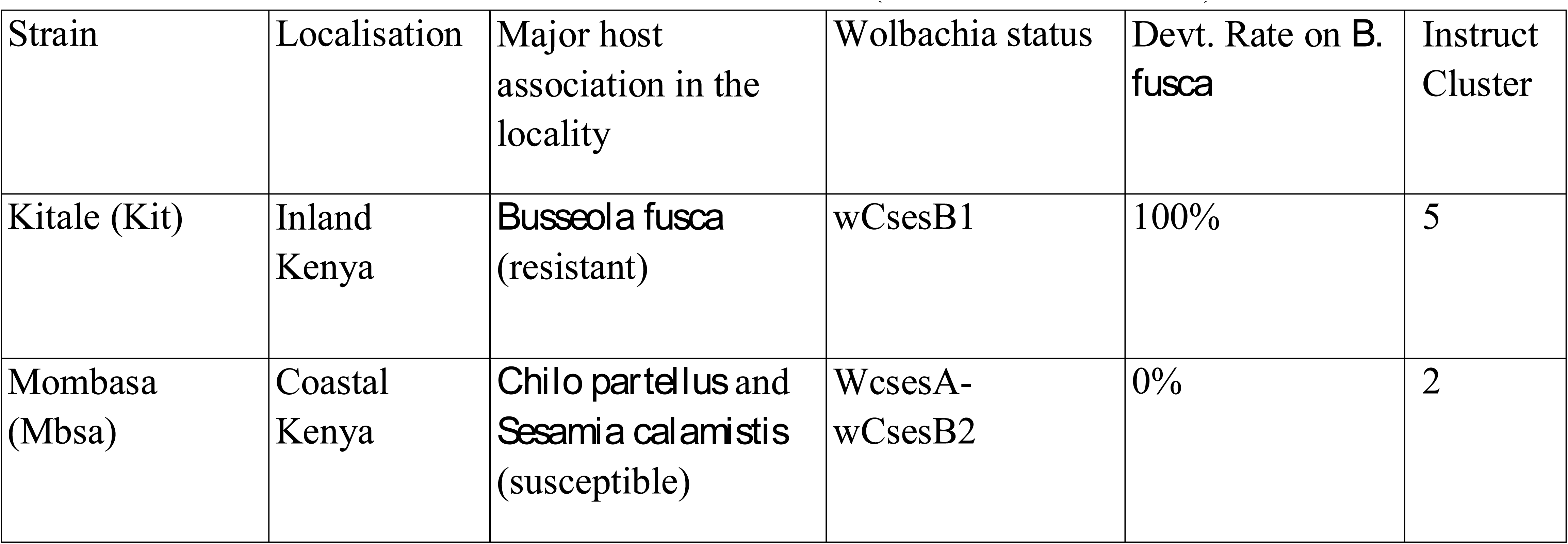
Status of the Kitale and Mombasa strains (Mochiah *et al*. 2002a)

| Strain | Localisation | Major host association in the locality | Wolbachia status | Devt. Rate on *B. fusca* | Instruct Cluster |
| --- | --- | --- | --- | --- | --- |
| Kitale (Kit) | Inland Kenya | *Busseola fusca* (resistant) | wCsesB1 | 100% | 5 |
| Mombasa (Mbsa) | Coastal Kenya | *Chilo partellus* and *Sesamia calamistis* (susceptible) | WcsesA-wCsesB2 | 0% | 2 |

Adults were fed a 20% honey–water solution imbibed in a cotton wool pad and kept under artificial light for 24 h to mate. In all experiments, only 1-day-old females, putatively mated and unexperienced to oviposition were used. The experiments were carried out at 25 ± 2 °C, 50– 80% RH, and a 12:12 h (L:D) photoperiod.

The stemborer species, *B. fusca* and *S. calamistis*, were continuously reared on artificial diet as previously Onyango & Ochieng’-Odero (1994). For each species, three times a year, several stemborer larvae were added to rejuvenate the colonies. Fourth larval instars were introduced into jars (10 x 20 cm), each containing pieces of maize stem, and left for 48 h to feed and produce frass to facilitate host acceptance by the parasitoid wasps for parasitism experiments. Thereafter, the larvae were used in the experiments.

### Genetic markers sequencing and genotyping

Eleven previously developed microsatellites markers (Jensen *et al.* 2002; Abercrombie *et al.* 2009) were amplified and fragment size determined. Amplifications were performed in 10 μL with approximately 5 ng of genomic DNA, 1 × HotStarTaq PCR buffer, 2 μL Q-Solution 5× (QIAGEN), 1.6 mM of dGTC, dTTP and dCTP, 50 μM dATP, 5 pM of each primer, 0.25 U Taq polymerase (HotStarTaq, QIAGEN) and 0.01 U of [33P]-dATP. The ‘touchdown’ PCR (Mellersh & Sampson 1993) was used as follows: initial activation step at 95°C for 15 min, 18 cycles at 94°C for 30 s, 60 to 51°C for 30 s (-0.5°C/cycle), 72°C for 30 s, 29 cycles at 94°C for 30 s, 54°C for 30 s, 72°C for 30 s and a final elongation step at 72°C for 10 min. Results were visualized using an ABI 310 and a AB3130 sequencer with fluorescent size standard (GeneScan 600 Liz, Applied Biosystem). Amplifications were made following conditions previously described using fluorescent labeling (Pet, Vic, Ned or 6Fam) of the forward primer.

Peaks identifying fragment sizes were assessed using GeneMapper 4.0 Software. Locus B1.42 presented peaks difficult to analyze with multiple bumps preventing any accurate measure of fragment size and was thereby discarded. Loci B1.155 and B5.126 were also not considered in the analyses because they presented a high percentage of missing genotypes (respectively 14.6% and 27.0%) probably reflecting the occurrence of null alleles. Eight loci were then genotyped per individual.

### Wolbachia

infection status was checked using the protocol developed in Branca *et al.* (2011).

### Cross-mating experiment

To obtain *Wolbachia*-free parasitoids colonies (named cured lines), the gravid females of each aforementioned parasitoid line were reared on larvae of *S. calamistis* previously fed on artificial diet Onyango & Ochieng’-Odero (1994), enriched with 2000 mg/L rifampicine (Dedeine *et al.* 2001). This process was repeated for three generations of female wasps to create cured colonies of Mombasa (Mbsa) and Kitale (Kit) *C. sesamiae*.

Cross experiment tests were conducted between both Mbsa and Kit *C. sesamiae* lines to assess the mating incompatibilities due to the presence of different *Wolbachia* types. Individual parasitoids were allowed to emerge singly by separating single cocoon from each cocoons mass. Individual male and female parasitoids from each colony (i.e. Kit *C. sesamiae* cured and uncured as well as Mbsa cured and uncured) were used for cross-mating experiments. Sixteen possible cross-mating combinations were investigated (Table 4). Each cross-mating combination was repeated at least 20 times.

After mating, females were presented 4th instar larvae of *S. calamistis* for oviposition using the method of Overholt *et al.* (1994). Thereafter, the larvae were reared and observed daily for mortality or parasitoid emergence. The developmental time of the progeny (egg to adult), the brood size, the sex ratio and the mortality outside and inside the host were recorded.

The presence of *Wolbachia* infections, in all *C. sesamiae* populations used in the cross-mating experiments, was tested using PCR techniques on *ftsZ* and *wsp* genes as described by Ngi-Song & Mochiah (2001). DNA was extracted from about 50 individuals (a mixture of males and females) from each population previously stored in 99% ethanol.

To test the effect of mating direction on each reproductive trait, a non-parametric Kruskal-Wallis test was applied with crosses as factor. ANOVA was not used because none of the data were normally distributed and had homoscedastic variance. Following Kruskal-Wallis test, a pairwise Wilcoxon’s rank sum test was conducted with false discovery rate (FDR) correction for multiple testing. Data were split into four groups for statistical analyses: crosses between Kit wasps, crosses between Mbsa wasps and crosses between populations in both directions. For all crosses, CI is expected between infected males and uninfected or differentially infected females. CI should lead to a reduction in female production either by female mortality (FM phenotype, diminution of the size of the progeny and the number of females) or male development (MD phenotype, only diminution of the proportion of females) (Vavre *et al.* 2000).

Statistical analyses for *Wolbachia* crosses experiments were performed in R 3.2 (R Core Team 2013).

### Genetic structure inference

To infer population structure from genetic data we used three different Bayesian methods for population partitioning: INSTRUCT, based on Hardy-Weinberg equilibrium with inbreeding (Gao *et al.* 2007), TESS3, taking into account spatial autocorrelation based on tessellation (Caye *et al.* 2016) and DAPC, a statistical partitioning method based on PCA (Jombart & Ahmed 2011). First, Instruct software was used with the Adaptive Independence Sample algorithm using inbreeding coefficient at population level as a prior model (mode 4, option v) (Gao *et al.* 2007), since *C. sesamiae* is known to have a highly inbred reproductive system (Ullyett 1935; Arakaki & Ganaha 1986). The number of clusters corresponding to the strongest genetic structure was determined using the method of Evanno *et al.* (2005). Each inference had a total number of iterations of 200,000 with a burn-in period of 100,000 iterations. Other parameters were kept as default value except the significance level of the posterior distribution of parameters, which was set to 0.95. The posterior probability of assignation of each individual was re-calculated over 10 MCMC runs using the CLUMPP software (Jakobsson & Rosenberg 2007) with greedy algorithm and 10,000 random permutations. Second, TESS3 was run using admixture with the BYM model (Durand et al. 2009). To identify the strongest structure, the model was run with K ranging from 2 to 9 using 100,000 sweeps with a 10,000 burning period. Degree of trend was assessed by running the algorithm with a varying value from 0 to 3 by 0.5 steps. The degree of trend showing the best DIC was kept. Genetic structure was then assessed for K=5, the best K, and T=1.5, the best degree of trend, with MCMC chain run for 1,000,000 sweeps with a 100,000 burn-in period. Third, we used a PCA-type approach with DAPC in R package *adegenet* (Jombart & Ahmed 2011) which is hypothesis-free since it just clusters individuals to maximize the explained genetic variance within the data.

The influence of various factors on the genetic variance was described using multiple correspondence analysis (MCA, package FactoMineR), and assessed using the *adonis* function in vegan R package (Oksanen *et al.* 2013). This corresponds to an extension of AMOVA (Excoffier *et al.* 1992) for crossed factors and in a non-hierarchical pattern (McArdle & Anderson 2001). The factors considered were: host genus, the *Wolbachia* infection status, spatial cluster of samples and the Köppen-Geiger climate type (Kottek *et al.* 2006). As the sampling was not done randomly across Sub-Saharan Africa, we tested the effect of spatial structuration by defining spatial cluster grouping localities close to each other. The spatial cluster of samples was obtained with hierarchical clustering from latitude and longitude data (*Mclust* function) (Fraley & Raftery 2002; Fraley *et al.* 2012). Genetic distance between individual were generated using Smouse Peakall’s formula (Smouse & Peakall 1999) in GenoDive (Meirmans & Van Tienderen 2004).

A Bayesian analysis of population sizes and reciprocal migration rates between the consensus genetic clusters obtained from partitioning methods was performed using software Migrate (Beerli & Palczewski 2010). Migrate-n software version 3.6.6 was run using the microsatellite model set to Brownian motion and the gene flow model to asymmetric. Since asymmetric gene flows can only be estimated pairwise, we run independently the software for each pairwise cluster comparison. Prior distributions of *θ* and *M* were chosen to get posterior distributions that are not truncated. Five chains of different heat from 1 to 10 were run for 500,000 generations with a burn-in period of 10,000.

## Results

### Genetic structure

The three clustering methods, Instruct, DAPC and TESS3 used in this study gave similar results regarding the population structure of *C. sesamiae* populations. For each method, the best fit was observed for five clusters (maximum delta-K for Instruct, Figure S1, diffNgroups criterion for DAPC and Deviance Information Criterion for TESS3). Regarding the structuration in relation to the host species, cluster 1 of all three methods was found exclusively on *Sesamia nonagrioides* (Figure 1, in red), clusters 2 and 3 were found mainly on *Busseola ssp.* (Figure 1, in yellow and green, respectively), cluster 4 on *Sesamia* and *Chilo spp.* (Figure 1, in blue) and finally cluster 5 was recovered from five host genera (Figure 1, in purple). Geographically, the three methods provided similar picture of genetic structure with some difference in admixture proportion. Cluster 1 population was scattered in between central Ethiopia, western Kenya and Northern Tanzania, and even Cameroon (Figure 2 in red). This corresponded to the population found on *S. nonagrioides.* One discordance appeared with the DAPC method, which failed to assign one individual from Arusha (Tanzania) into Cluster 1. Cluster 2 extended from Eritrea to Western Kenya in Instruct, but was restricted to Western Kenya in TESS3 and DAPC (Figure 2, yellow). Conversely, cluster 3 was only present in Western Kenya and central Tanzania in the three methods but extended to Eritrea in TESS3 and DAPC (Figure 2, green). Cluster 4 extended from South Africa to East Kenya and Rwanda in the three methods (Figure 2, blue). In Instruct and TESS3 analyses, a very high posterior probability of cluster 4 was also observed further west in the coast of Congo-Brazzaville. Cluster 5 extended from Tanzania to Cameroon in all three methods but was found much more spread in DAPC analysis, until South-Africa, and to a lesser extent in Instruct (Figure 2, purple). Overall, there seem to be a clear delimitation between cluster 2 and 3, which extend form Tanzania to Eritrea, and cluster 4 and 5, which were found from Cameroon to South Africa. Delimitation within these two groups of two clusters seemed to be shallower and influenced by the method used.

**Figure 1.**
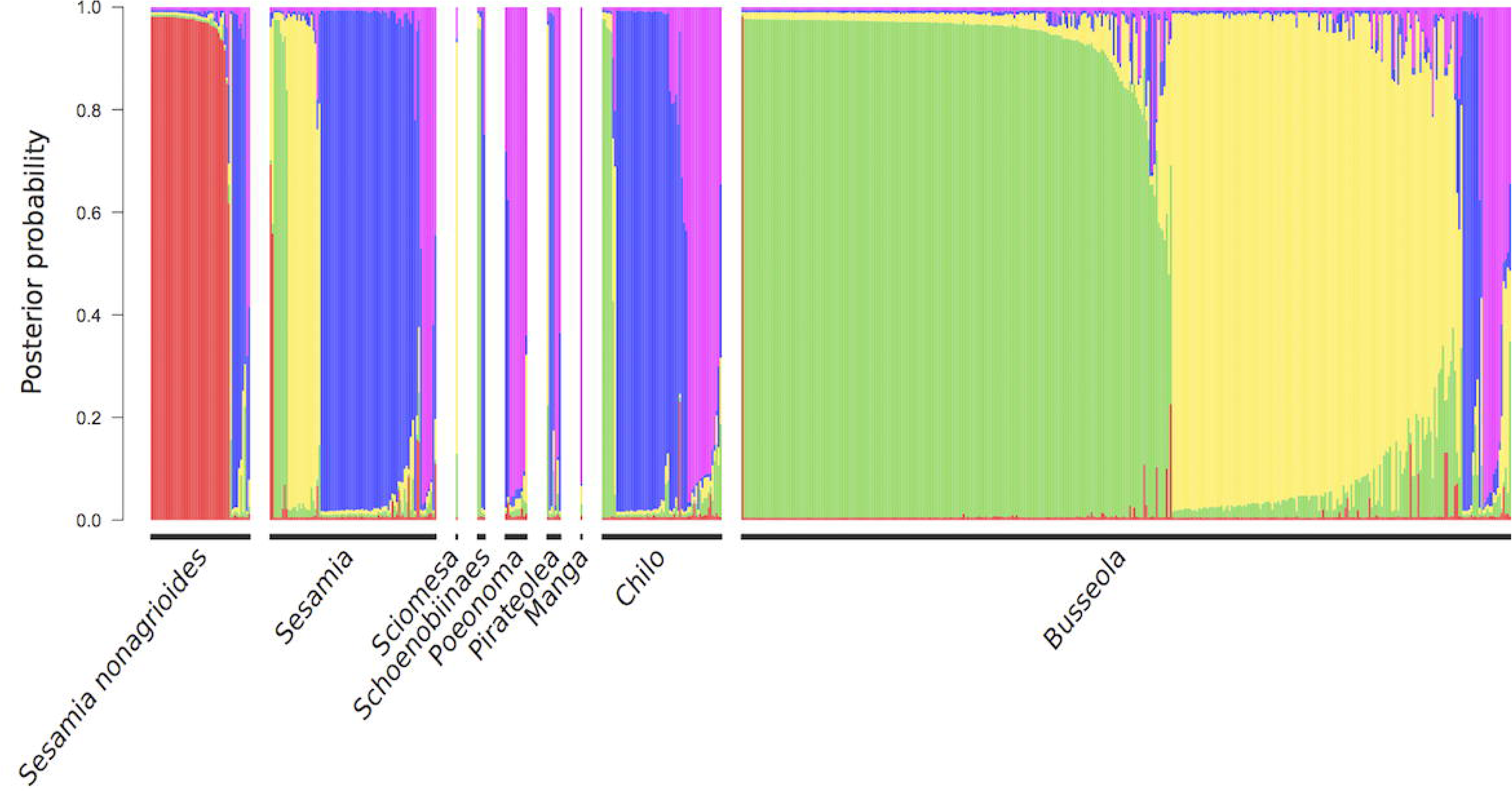
Posterior probability of assignment of each individual of *Cotesia sesamiae* wasps to each of the 5 Instruct clusters and post-processed with CLUMPP (Cluster 1 in red, cluster 2 in yellow, cluster 3 in green, cluster 4 in blue, cluster 5 in purple). Individuals are grouped by the host genus where they were found. Individuals found on an unidentified host are not represented.

**Figure 2.**
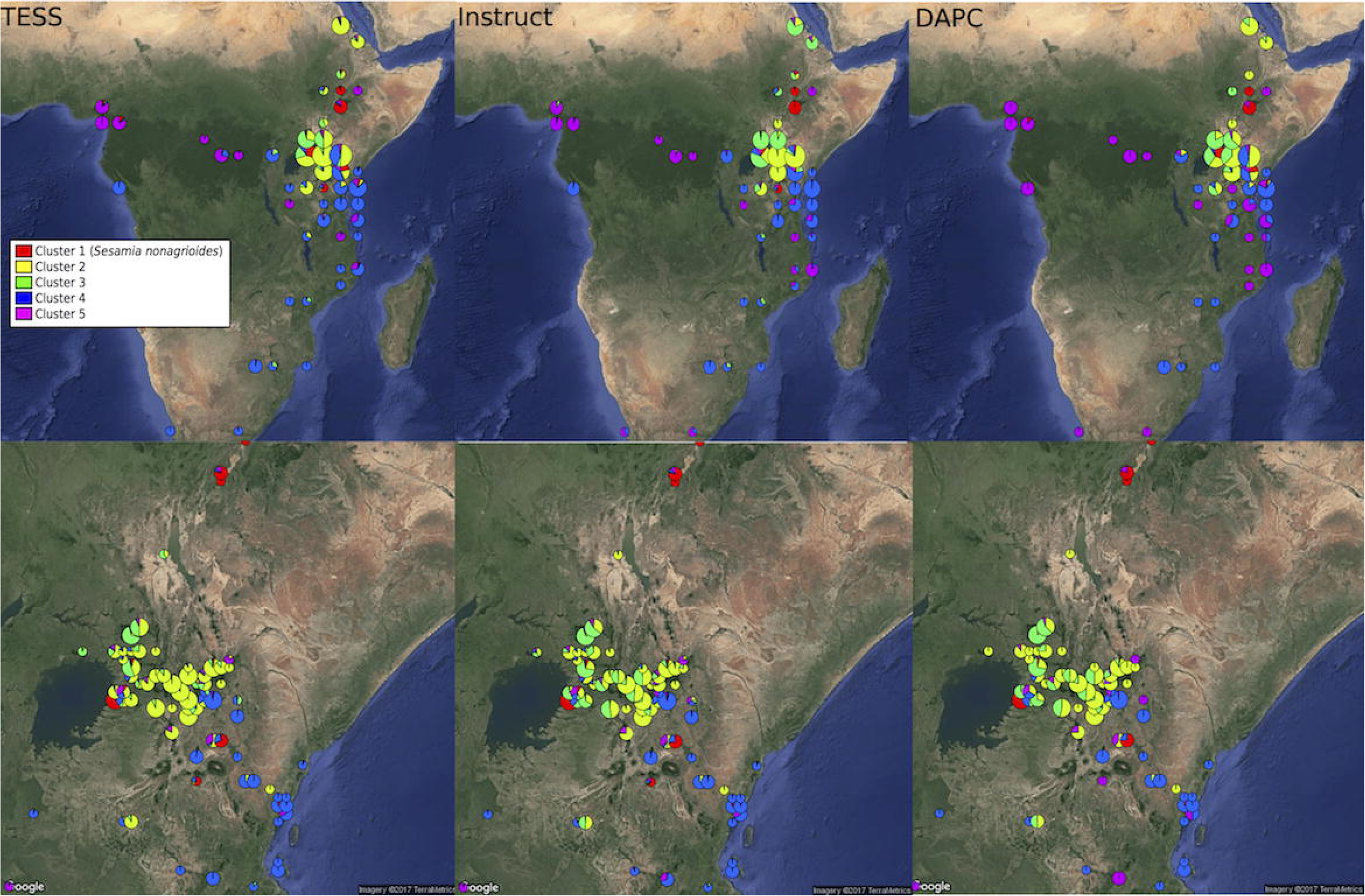
Distribution of genetic clusters of *Cotesia sesamiae* wasps for DAPC with K=5 (A), TESS3 software (B) and the Instruct software CLUMPP consensus with K=5 (C). For each clustering method, only individual with posterior probability of assignment above 0.5 are represented for each analysis. Distribution in Sub Saharan Africa is represented at the top and a zoom in Kenya at the bottom.

### Wolbachia *strains distributions*

A rather good concordance was observed between genetic structure at microsatellites level and *Wolbachia* strain distributions (Figure 3). Cluster 1 was associated exclusively with the *Wolbachia* wCsesA strain, cluster 2 and 3 with wCsesB1, cluster 4 and 5, with the bi-infection wCsesA/wCsesB2.

**Figure 3.**
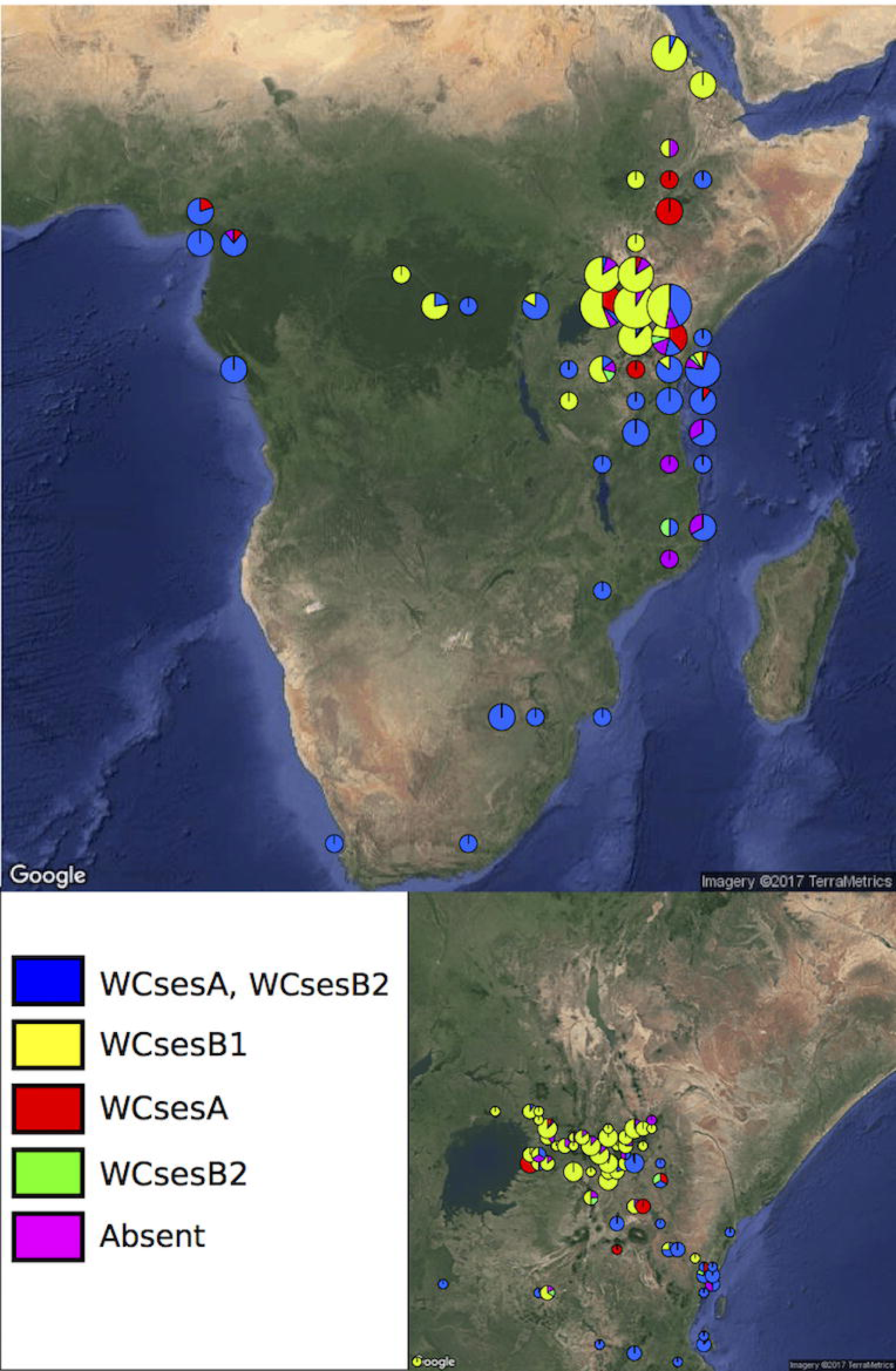
Distribution of *Wolbachia* infection in *Cotesia sesamiae* wasps across Sub-Saharan Africa. Red: wCsesA,: blue: wCsesA/wCsesB2, yellow: Absent, purple: wCsesB2, green: wCsesB1.

### Relative influence of biotic and abiotic factors

The individuals belonging to the cluster found exclusively on *Sesamia nonagrioides* were interpreted as a distinct species by Kaiser *et al.* (2015, 2017), based on eco-phylogeny analyses and cross-mating experiments, and corresponding to a host and plant-host driven ecological speciation event. As it has now been described as the species *Cotesia typhae* (Kaiser et al. 2017), it was not considered in these analyses to prevent an overestimation of host effect. Multiple correspondence analysis (Figure 4) suggested the presence of structure in relation to all the factors considered (spatial cluster, Köppen-Geiger climate classification, *Wolbachia* infection status and host genus). The full models tested with *adonis* function (Table 2) confirmed that all neutral (geography) and selective forces, abiotic (climate type and geography) and biotic (host genus, *Wolbachia* infection), contribute significantly to the genetic variance of the microsatellite genotypes. Because the *adonis* method tests factors sequentially, it is important to consider each factor as either first term or marginal (last) term to see the extent of the effect. When added first in the sequence of factors in the *adonis* function, the biotic factors had higher R^2^ than the abiotic factors (0.43 and 0.38 for *Wolbachia* and host genus, respectively and 0.28 and 0.21 for Köppen-Geiger Climate type, and localization, respectively) (Table 3). In addition, all the factors had significant marginal effects (Table 3). Pairwise interactions between factors were weak (R^2^<0.04), but significant for all the possible interactions (Table 2). None of the tripartite interactions was significant.

**Table 2.**
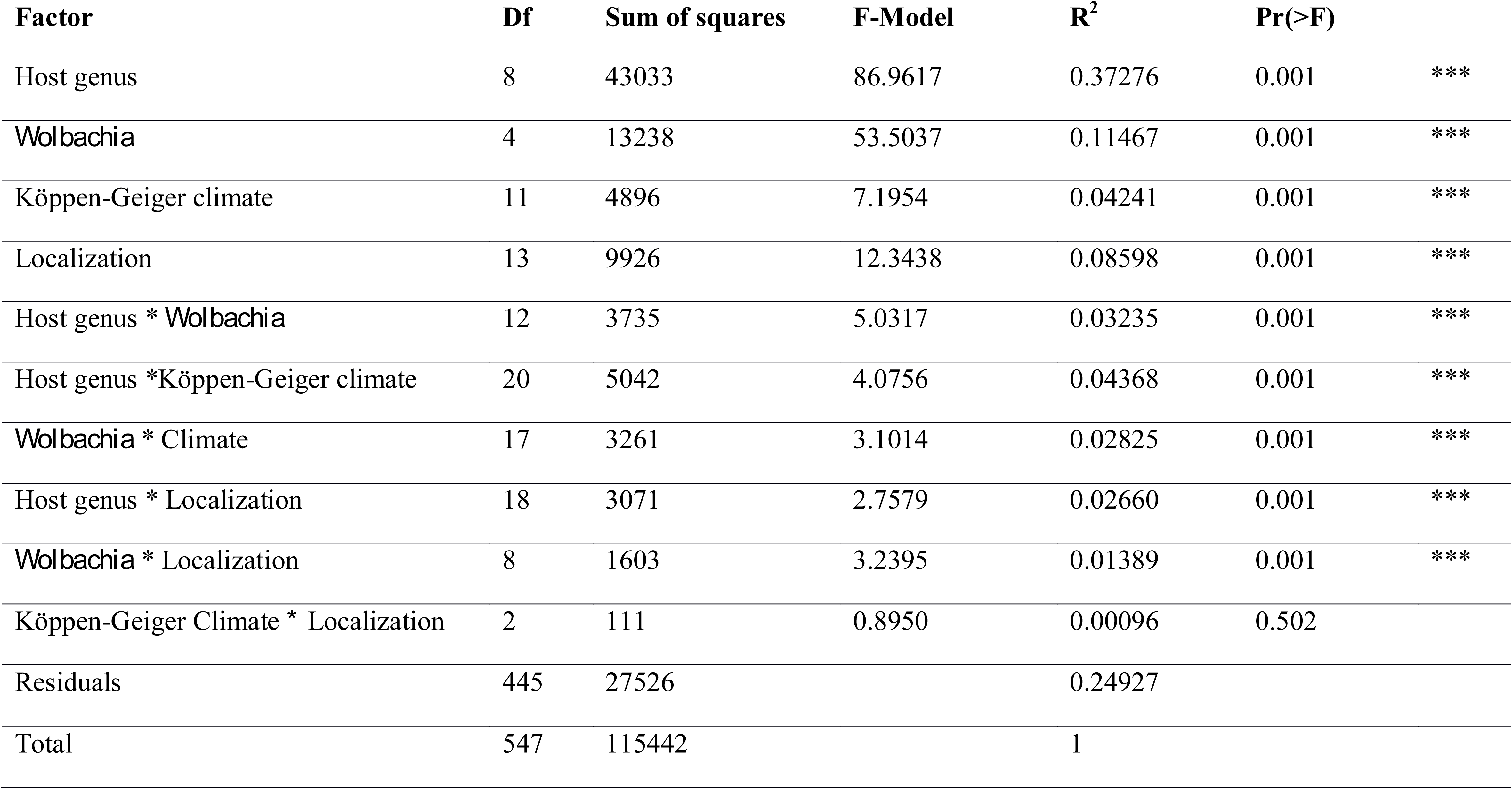
Analysis of molecular variance using microsatellite distance matrices and a full model containing all terms and interactions.

**Table 3.** Sum of squares and partial R^2^ of Host genus, *Wolbachia* infection status, Köppen-Geiger climate and localization taken either as marginal effect or as the first term when adding them sequentially.

| **Factor** | **Df** | **Marginal**<br>**Sum of squares** | **Marginal**<br>**Partial $R^2$** | **1<sup>st</sup> Sequential**<br>**Sum of squares** | **1<sup>st</sup> Sequential**<br>**Partial $R^2$** |
| --- | --- | --- | --- | --- | --- |
| Host genus | 8 | 3709 | 0.03213 | 43033 | 0.37277 |
| *Wolbachia* | 4 | 5711 | 0.04947 | 49576 | 0.42944 |
| Köppen-Geiger climate | 11 | 2473 | 0.02142 | 32467 | 0.28124 |
| Localization | 13 | 9926 | 0.08598 | 34013 | 0.29463 |

**Table 4.**
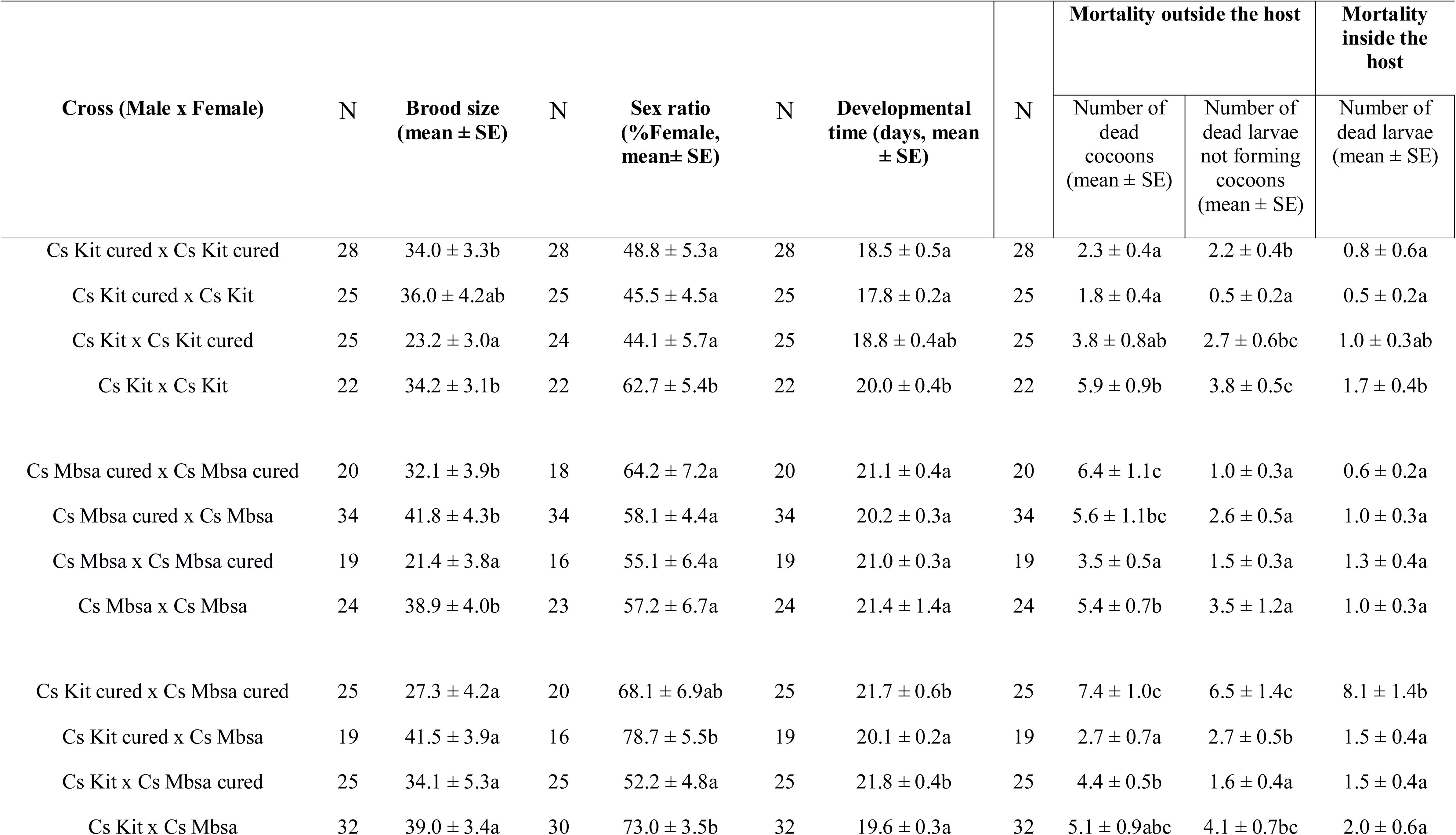
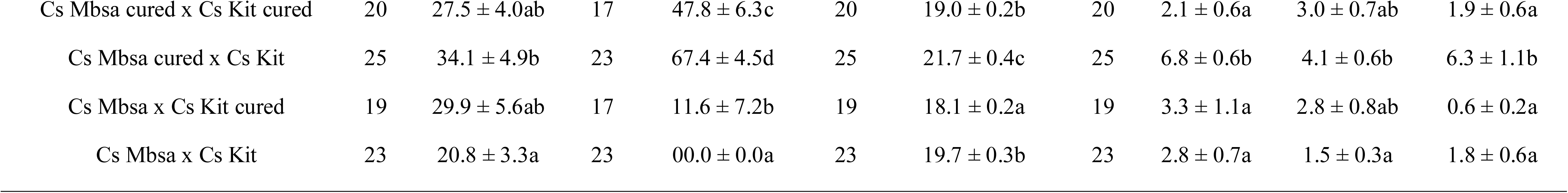
Brood size, sex ratio, developmental time and mortality outside and inside the host of populations of different crosses on *Sesamia calamistis* (N = number of replicates). Note. Cs Kit, *Cotesia sesamiae* from the inland Kitale area of Kenya; Cs Mbsa, *Cotesia sesamiae* from the coastal Mombasa area of Kenya; cured, *Wolbachia*-free parasitoids colonies (i.e. cured lines); in crosses within each population and between populations, values with different letter are significant (q-value <0.05; pairwise Wilcoxon’s rank sum test, q-value = FDR corrected p-value).

**Figure 4.**
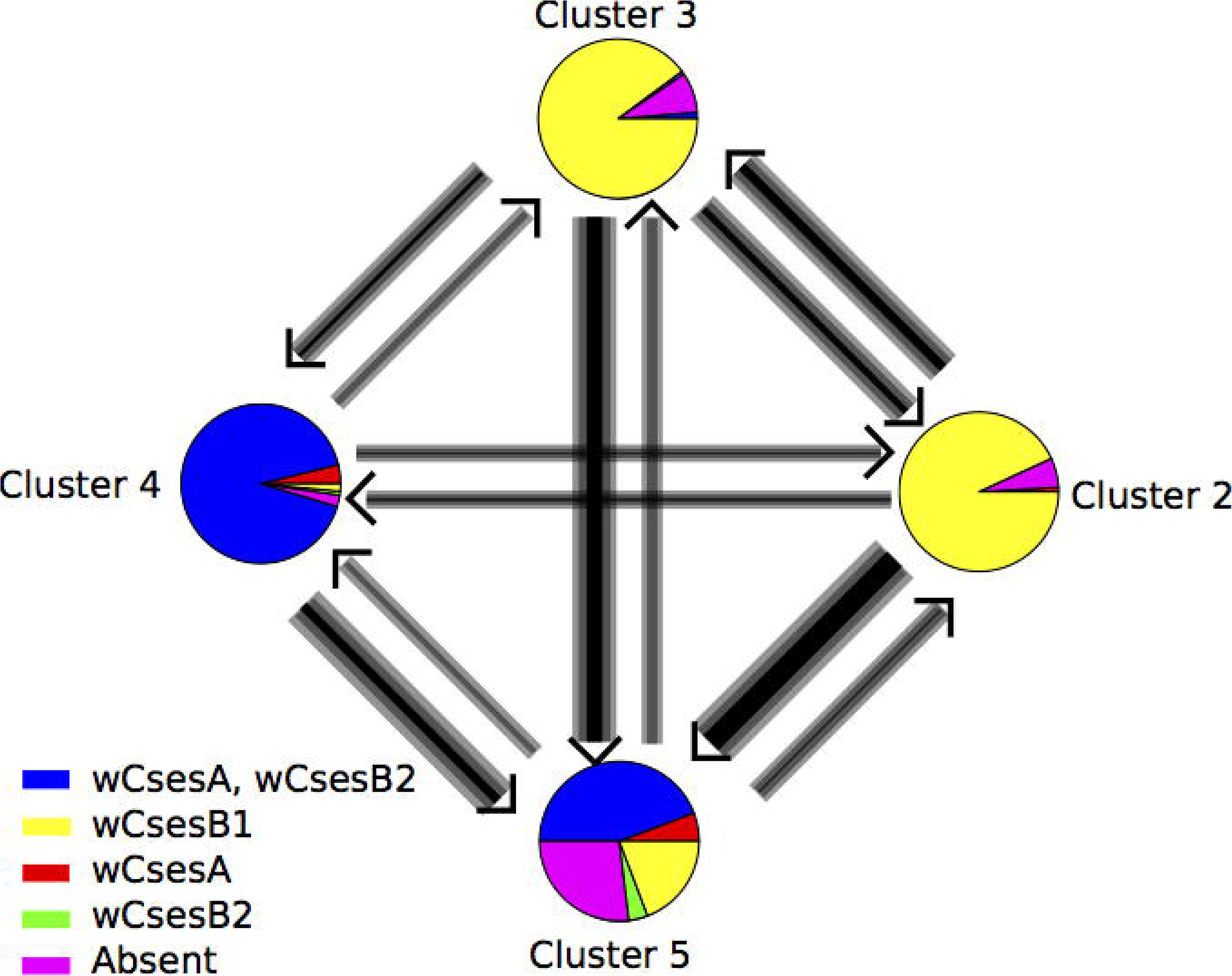
Estimates of gene flow between four geographic genetic clusters of *Cotesia sesamiae* wasps identified by Instruct. Each circle represents the infection status of individual found assigned to each cluster and the colour are corresponding to the one on figure 3. The fifth genetic cluster found only infecting *Sesamia nonagrioides* was excluded of the analysis.

### Wolbachia *crosses experiment*

For crosses within each population, the brood size dropped in crosses involving infected males and cured females (i.e. Cs Kit x Cs Kit-cured and Cs Mbsa x Cs Mbsa-cured) from 34-36 to 23 for Kitale population and from 32-42 to 21 for Mombasa population (Table 4). Although for these both potentially incompatible crosses the sex ratio (or %females) decreased significantly for Kitale population and no significant change was detected for Mbsa population, the overall number of females was reduced in these both crosses from 45-62% to 44% and from 57-64% to 55% for Kitale and Mbsa population, respectively. No significant changes in the developmental time and in the mortalities outside and inside the host through dissection were detected between these incompatible crosses and the other crosses.

To the contrary, in crosses potentially showing bidirectional CI, i.e. crosses involving individuals from different populations and infected with different *Wolbachia* strains (i.e. Cs Kit x Cs Mbsa and Cs Mbsa x Cs Kit), we only found a significant decrease of the percentage of female from 47-67% to 11-0% in the cross involving Mbsa males and Kit females (Table 4). In this cross, almost no females were recovered despite a normal overall progeny size, suggesting a complete incompatibility with pure male development (MD) phenotype (Vavre *et al.* 2000). By contrast, in the cross for Kit male (wCsesB1) with Mbsa female (wCsesA/wCsesB2), CI expressed only when Mbsa females were cured and only partially since females were recovered. No significant changes in the developmental time and in the mortalities outside and inside the host through dissection were detected between these incompatible crosses (i.e. Cs Kit x Cs Mbsa-cured, Cs Kit x Cs Mbsa, Cs Mbsa x Cs Kit-cured and Cs Mbsa x Cs Kit) and the other crosses.

### Migration patterns

For Bayesian analyses of pairwise migration rates, acceptance rate ranged between 0.20 and 0.56 with an effective MCMC sample size from ∼500 to ∼2700. Clusters defined by Instruct were used except Cluster 1 for the main reasons as exposed above. Mostly symmetric gene flow was found between Cluster 2 and 3, which are mainly infected with the same wCsesB1 *Wolbachia* strain (Figure 5); they were found mainly on *Busseola,* at least in one contact zone in Central Kenya (Figure 2). Otherwise, asymmetric gene flow between clusters were found. All the gene flow with cluster 5 were orientated toward Cluster 5. Gene flow between Cluster 4 (found mainly on *Sesamia* and *Chilo*) and Cluster 2 was the lowest despite the presence of a contact zone in Kenya (Figure 2). Kit population from the laboratory colony was assigned to Cluster 2 and Mbsa population from the laboratory rearing to cluster 5 as inferred in Instruct clustering (Table 1). Therefore, migration was more orientated from wCsesB1-infected population toward wCsesA/wCsesB2 bi-infected populations, mainly because of an asymmetric gene flow in that particular direction between Cluster 3 and Cluster 4.

**Figure 5.**
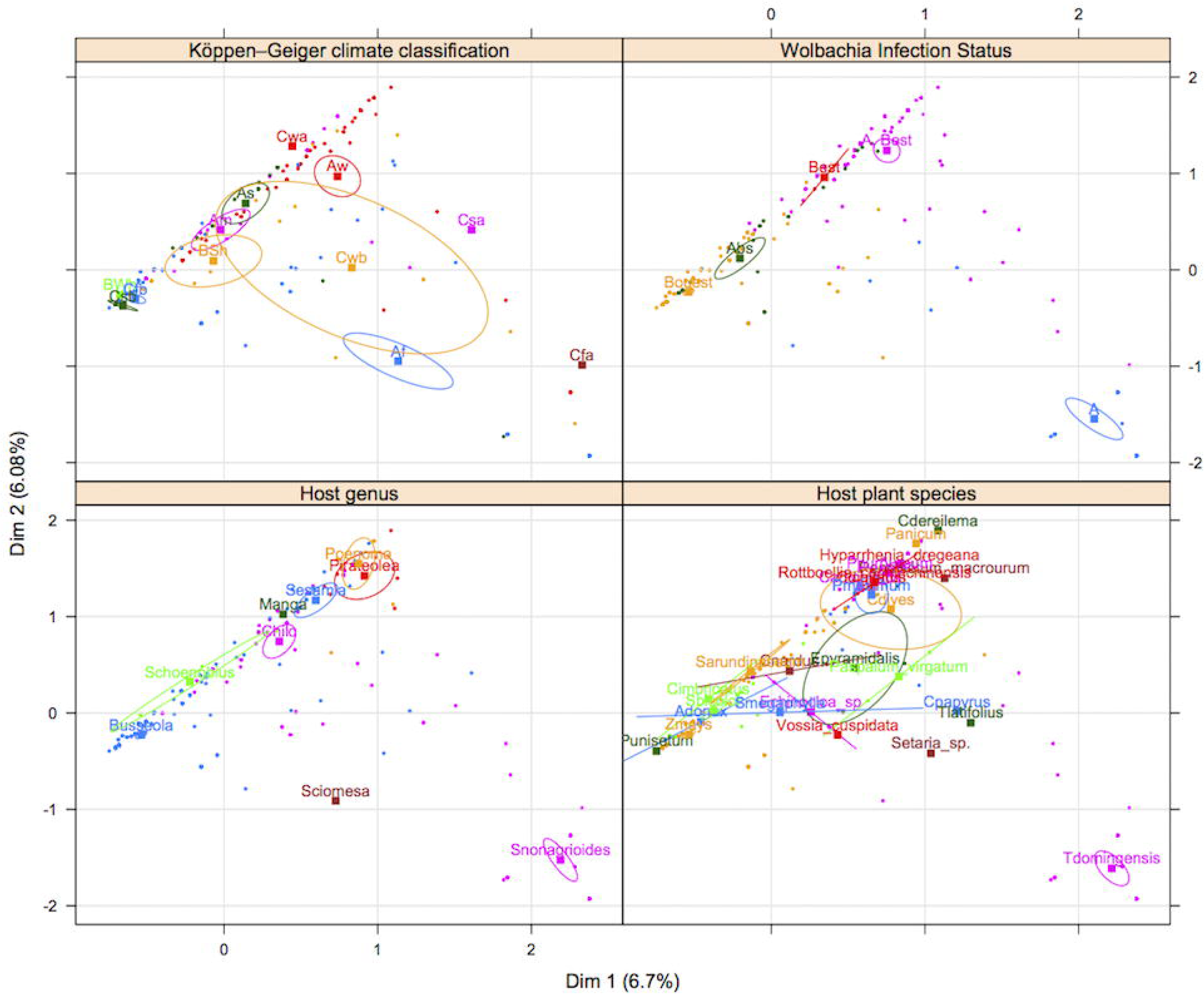
Multiple correspondence analysis between microsatellite markers distance between individuals and ecological variables

## Discussion

### *Geographic, ecological and biotic factors determining the genetic structure of* Cotesia sesamiae

The five major clusters inferred by the three different genetic clustering methods, TESS, Instruct and DAPC exhibited very similar geographic partition. However, TESS3 and Instruct admixture models were more concordant. DAPC results differed by the many geographic areas assigned to just one cluster. The DAPC algorithm optimizes a model without admixture that assigns individuals and not portion of their genomes to clusters. It seeks linear combination of genetic variables that maximizes between clusters component of the genetic distance between individuals. Models without admixture are not robust to the inclusion of admixed individual in the sample, reciprocally, models with admixture are less able to detect barriers when admixture is limited (François *et al.* 2010). In the absence of intrinsic biological reproductive barriers between the populations, we would expect the admixture model is the best suited because the five clusters are all represented in Kenya and Tanzania with a geographic continuum in both countries. However, the presence of reproductive isolation mechanisms, may limit admixture in this continuum of populations. Indeed, the results of Instruct non-spatial admixture model (Figure 2) shows that populations maintained their integrity; admixture being limited to the hybridization zones despite the ability of *C. sesamiae* to expand throughout Africa. We will discuss below the factors that may limit admixture in this continuous environment in the light of our results on experimental crosses, *Wolbachia* strains distribution, host ecological specialization, climate, and on the biology of *C. sesamiae*.

There are at least three strains of *Wolbachia* infecting *C. sesamiae* populations across Sub-Saharan Africa (Branca *et al.* 2011). We did not find bidirectional incompatibility between populations as a result of different infection. Only individuals infected with wCsesA and wCsesB2 strains showed incompatibility with cured or wCsesB1 infected Kit individual. Infected individuals wCsesA/wCsesB2 were already found highly incompatible in a previous study (Mochiah *et al.* 2002b), but incompatibility was not assessed for wCsesB1-infected individuals. The results for *Wolbachia* crosses involving wCsesB1 infected males and cured females does not present normal CI phenotype because there was no increase in male proportion in the progeny; however, we observed a reduction in progeny size (males and females) leading to a reduced number of females. This result is coherent with *Wolbachia* invasion theory since *Wolbachia* fitness is linked to the fitness, which female progeny size is a proxy, of *Wolbachia*-infected females relative to non-infected counterparts (Werren, 1997). However, this means that there is an unknown mechanism leading to high mortality of male eggs in incompatible crosses. Possibly, part of the male progeny includes in fact diploid males that could be affected in incompatible crosses, as diploid males are common in *Cotesia* wasps (Zhou *et al.* 2006; De Boer *et al.* 2007). A direct effect on development, not related to fertilization, could be also considered. In a similar way, surprisingly, no strong incompatibility was observed between Mbsa wCsesA/wCsesB2 cured female and Mbsa infected males as no biased sex ratio was found in the progeny. However, as in the case of Kit, a reduction in progeny size was observed which means that probably CI expresses differently between individual of the same genetic background (Kit or Mbsa), than when incompatible crosses occur between different genetic backgrounds. In the inter-population crosses studied here, a MD phenotype is very likely, as male-biased sex ratio was not associated to significant progeny size reduction. The consequence of this unidirectional incompatibility will be asymmetric gene flow between differentially infected populations (Jaenike *et al.* 2006; Telschow *et al.* 2006). Indeed, CI is an efficient mechanism for *Wolbachia* to spread within populations by giving infected females a higher fitness. We should therefore expect the spread of individuals infected with wCsesA/wCsesB2 across *C. sesamiae* geographical range, reflected by higher migration rate from wCsesA/wCsesB2-infected populations toward other populations. However, using microsatellite markers, we observed conversely a lower migration rate from wCsesA/wCsesB2 -infected genetic clusters toward the other clusters (Figure 5), except for the migration between cluster 4 and 2. This unexpected result may be explained by local adaptation. Regions where *C. sesamiae* are infected with wCsesA/wCsesB2 are indeed dominated by avirulent parasitoids and susceptible hosts, whereas regions where *C. sesamiae* are infected with wCsesB1 are dominated by virulent parasitoid attacking resistant host. Females migrating from bi-infected to wCsesB1 regions are maladapted and killed by encapsulation, but females migrating from wCsesB1 to regions with wCsesA/wCsesB2-infected individuals are able to develop on the host. Yet, males infected with wCsesB1 can reproduce with bi-infected females wCsesA/wCsesB2, which would allow some gene flow from wCsesB1 to wCsesA/wCsesB2. In conclusion, *Wolbachia* incompatibility favors the expansion of avirulent parasitoid wasp that are not capable to survive in some areas, and, in the opposite, the spread of virulent parasitoid is limited by area where parasitoid population are dominated by individuals infected with highly incompatible Wolbachia. This situation should lead to potentially stable contact zone between populations and current genetic structure.

To disentangle the effects of geography, *Wolbachia* infection, parasitoid host, and other ecological factors, a statistical model was optimized using *adonis* R function. The biotic and abiotic factors including geography analyzed in our statistical model explained more than 75% of the genetic variance. When looking at the factors most correlated to the genetic structure, our results are consistent with the hypothesis that ecology plays a significant role in reinforcing *C. sesamiae* population structure across evolutionary time. Indeed, *adonis* analysis showed that the strongest determinant of genetic variance was *Wolbachia* infection followed by the host species and the least contributing factors were localization and climate. An illustration of the dominant effect of the host is the particular status of the population represented by cluster 1, consistently collected on *Sesamia nonagrioides*. This population shows also higher Fst when compared to the other populations in every clustering method confirming it constitutes a new species as it has recently been proposed (Kaiser *et al.* 2015, 2017). Another population corresponding to cluster 5 expands from Cameroon to East-Africa and Uganda, trough Democratic Republic of Congo; this region corresponds to the great Equatorial forest of Africa, which is characterized by hot and wet climatic conditions. The cluster 4, located from Eastern Kenya to Mozambique along the Coast, is situated in a much drier area than cluster 5. This area is also important regarding host, since *B. fusca*, characterized as a resistant host, is rare in those regions (Le Ru *et al.* 2006; Moolman *et al.* 2014). The cluster 2 and 3 are located both in North-Eastern Sub-Saharan Africa but their positions differed according to clustering algorithm (i.e. West Kenya, Ethiopia and Eritrea). In terms of climatic conditions, these regions are very similar but the observed clusters might reflect two sympatric populations with recurrent gene flow as they are infected with the same *Wolbachia* strains (Figure 3).

These *C. sesamiae* populations show some geographic similarities with the genetic structure observed in the known resistant host *B. fusca* (Dupas *et al*. 2014), with five clusters observed across Africa, and a strong structure observed in East African Rift Valleys regions, contrasting with reduced structure observed in South and Central African regions. The cluster 3 of *C. sesamiae* located between Eastern and Western Rift Valley has an overlapping distribution with “H” cluster of *B. fusca*. The cluster 2 on the East of Eastern Rift overlaps with “KE” cluster of *B. fusca*. The cluster 4 of *C. sesamiae* ranges in East Africa at lower altitudes where *B. fusca* is rare or absent (Dupas *et al*. 2014) and to the south. The clusters 4 and 5 exhibit large distributions that overlap with the “S” cluster of *B. fusca* from South to East and Central Africa (Figure 2). A fifth population is also present in both species. Cluster 1 of *C. sesamiae* corresponds to parasitic wasp infecting *S. nonagrioides* that has been described as a new species, *C. typhae*. Cluster “W” of *B. fusca* is only present in West Africa and isolated from the other *B. fusca* populations (Figure 2). These results suggest that *B. fusca* and *C. sesamiae* share a common phylogeographic history that explain the current genetic structure of both species. For instance, the highest diversity for both species has been found in the East African Rift Valley. The East African Rift valley also explained the differentiation observed between two *C. sesamiae* lineages based on 6 mitochondrial and nuclear markers (Kaiser *et al.*,2015). One lineage corresponds geographically and ecologically to clusters 2 and 3, and the second one to cluster 4. The East African Rift Valley has already been observed as a center of diversification for several species (Odee *et al.,* 2012; Habel et al, 2015; Freilich et al, 2016; Mairal *et al.*, 2017). This observed biological diversity has been related to both topological heterogeneity and variable climatic conditions that occurred since the formation of the East African Rift Valley ca. 20 Mya, with the alternation of arid and wet periods (Sepulchre *et al*., 2006). Therefore, we could explain this observed pattern either by first the colonization of the East African Rift Valley followed by diversification or that the origin of both species lays in the East African East Valley which has been followed by further extension with admixture across Africa, except in West Africa, where *C. sesamiae* is absent and where *B. fusca* is totally isolated with zero migration observed to date (Sezonlin *et al.* 2006; Dupas *et al.* 2014).

### Wolbachia *and biological control*

It is widely acknowledged that a better understanding of tritrophic interactions between plants, phytophagous insects and associated antagonists can help to develop better pest management strategies by identifying bottom-up and top-down effects in the food chain (Agrawal, 2000). *Wolbachia* can be considered as a fourth trophic level in such system but, the impact of *Wolbachia* on parasitoid host plant interactions has not received much attention. It was shown that a *Wolbachia* strain invasion temporarily reduces the impact of the parasitoid on its host (Branca & Dupas 2006). But this impact can be sustained in the case of stable contact between incompatible strains in “hybrid” zones. Conversely, *Wolbachia* can reinforce adaptive divergence between locally adapted populations to the benefit of the parasitoid (Branca *et al.* 2009). *Cotesia sesamiae* is a good model to test the effect of *Wolbachia* on host parasitoid assemblages as the four consensus genetic clusters differed for their *Wolbachia* and Lepidoptera host associations. In hybrid area, maladaptive gene flow may be observed and limited by *Wolbachia* strain bidirectional incompatibility. This is the case between coastal (Mbsa) and inland (Kit) populations of the parasitoid (Dupas *et al.* 2008). The maladaptation may be the strongest in the AS Köppen Geiger Climate Zone (corresponding to dry mid altitude agroclimatic zone) in wet seasons when *B. fusca* represents half of the host community (Ong’amo *et al.* 2006), whereas avirulent *C. sesamiae* toward *B. fusca* dominates parasitoid populations (Dupas *et al.* 2008). Strong counter counter-selection of avirulent alleles is expected in *B. fusca* abundance peaks. *Busseola fusca* is dominant in some seasons in mid altitude areas where virulent alleles dominate (Dupas *et al.* 2008). Although avirulent parasitoids are able to select host at contact, which may reduce maladaptation in the field, using contact cues to select host is risky because the host can bite and kill the parasitoid before oviposition can be made; 25% of *C. sesamiae* entering the stem tunnel are killed by *S. calamistis* larvae upon contact (Potting *et al.* 1999). The presence of partially incompatible *Wolbachia* strains in the virulent and avirulent parasitoid populations may favor their cohesiveness in balancing host communities across seasons. Hence, reducing gene flow between locally adapted populations toward their host, in absence of premating isolation, might reduce maladaptation in hybrid zone and our study confirms *Wolbachia* can reinforce this process. For instance, very few heterozygous females between virulent and avirulent alleles on the bracovirus CrV1 locus has been found in a previous study, since they are likely maladapted (Branca *et al.*, 2011). Therefore, we would expect a lack of recombination and strong diversification on genes, particularly at the bracovirus locus, related to host specificity in *C. sesamiae*, pattern that has yet to be investigated at the genome level.

Thompson (2005), in his seminal book on coevolutionary mosaics stressed that gene flow had an ambivalent influence on coevolutionary interactions. Gene flow is essentially maladaptive, bringing locally maladapted genes to populations in interaction (Nuismer 2006), but in the presence of negative frequency dependent dynamics of coevolutionary interactions, rare new variants originating from other populations may be adapted. Our results show some congruence between *C. sesamiae* and *B. fusca* genetic structure (Dupas *et al.* 2014). Congruence with host structure is therefore observed at different ecological levels, not only at the level of host genus as shown by *adonis* analyses but also at the level of host populations. This may reduce maladaptation of *C. sesamiae* toward *B. fusca* and favor local coevolutionary interactions.

## Conclusion

Our study presents a unique comprehensive case for assessing the determinant of genetic structure in a parasitoid species, including multiple interactive biotic and abiotic forces. The parasitoid, like its main host *B. fusca*, likely diversified across the East African Rift Valleys where all the genetic clusters are found. Despite their wide distribution across Sub-Saharan Africa, some populations have maintained their integrity as shown by the non-spatial admixture model. Two important results pinpoint toward the strong influence of host on parasitoids population dynamics and population genetics at a large geographical scale: (1) although the species genetic clusters appear to have diversified across East African Rift Valleys refuges, host species that are distributed across Africa became then the strongest factor determining genetic structure, rather than climatic selection and geographic isolation (2) migration rate inferred from Bayesian analysis of microsatellite data suggests a limitation of gene flow due more to host adaptation than to *Wolbachia* infections. This result has a fundamental importance in the context of biological control program. As opposed to chemical control agents, biological control agents are expected to be able to cope with host evolution (Holt & Hochberg 1997) but other interactions may limit this evolutionary sustainability. In our case, parasitoid wasps are able to cope with host evolution despite many additional biotic and abiotic ecological forces including reproduction manipulators that would be expected to reduce local adaptation to host. The insect host dominates the piling up of all these factors and could explain why parasitoids can be very successful biological control agents even when introduced in climatically and geographically distant environments from their native settings (Stiling & Cornelissen 2005). More generally, this work supports the hypothesis of the higher impact of ecological *versus* neutral forces and of host *versus* other ecological forces on the diversification of parasitoid - host interactions.

## References

Abercrombie, L. G., C. M. Anderson, B. G. Baldwin, I.C. Bang, R. Beldade, G. Bernardi, A. Boubou, et al. 2009. « Permanent genetic resources added to Molecular Ecology Resources database 1 January 2009–30 April 2009 ». Molecular Ecology Resources 9 (5): 1375–1379.

Agrawal, A. A. 2000. « Mechanisms, ecological consequences and agricultural implications of tri-trophic interactions ». Current Opinion in Plant Biology 3 (4): 329–35

Antolin, M. F., & D. R. Strong. 1987. « Long-distance dispersal by a parasitoid (Anagrus delicatus, Mymaridae) and its host ». Oecologia 73 (2): 288–292.

Arakaki, N., & Y. Ganaha. 1986. « Emergence pattern and mating behavior of Apanteles flavipes (Cameron)(Hymenoptera: Braconidae) ». Applied Entomology and Zoology (Japan).

Asgari, S, O Schmidt, & U. Theopold. 1997. « A polydnavirus-encoded protein of an endoparasitoid wasp is an immune suppressor ». J Gen Virol 78 (11): 3061–70.

Assefa, Y., A. Mitchell, D. E. Conlong, & K. A. Muirhead. 2008. « Establishment of Cotesia flavipes (Hymenoptera: Braconidae) in sugarcane fields of Ethiopia and origin of founding population ». Journal of economic entomology 101 (3): 686–691.

Beckage, N. E.. 1998. « Modulation of immune responses to parasitoids by polydnaviruses ». Parasitology 116: 57–64.

Beerli, P., & M. Palczewski. 2010. « Unified Framework to Evaluate Panmixia and Migration Direction Among Multiple Sampling Locations ».

Bordenstein, S. R., F. P. O’Hara, & J. H. Werren. 2001. « Wolbachia-induced incompatibility precedes other hybrid incompatibilities in Nasonia ». Nature.

Branca, A., & S. Dupas. 2006. « A model for the study of Wolbachia pipientis Hertig (Rickettsiales: Rickettsiaceae)-induced cytoplasmic incompatibility in arrhenotokous haplodiploid populations: consequences for biological control ». Annales de la Société Entomologique de France 42 (3-4): 443–48.

Branca, A., B. P. Le Ru, F. Vavre, J. F. Silvain, & S. Dupas. 2011. « Intraspecific specialization of the generalist parasitoid Cotesia sesamiae revealed by polyDNAvirus polymorphism and associated with different Wolbachia infection. » Molecular Ecology.

Branca, A., F. Vavre, J. F Silvain, & S. Dupas. 2009. « Maintenance of adaptive differentiation by Wolbachia induced bidirectional cytoplasmic incompatibility: the importance of sib-mating and genetic systems ». BMC Evolutionary Biology 9 (1): 185.

De Boer, J. G., P. J. Ode, L. E. M. Vet, J. B. Whitfield, & G. E. Heimpel. 2007. « Diploid males sire triploid daughters and sons in the parasitoid wasp Cotesia vestalis ». Heredity 99 (3): 288–294.

Dedeine, F., F. Vavre, F. Fleury, B. Loppin, M. E. Hochberg, & M. Boulétreau. 2001. « Removing symbiotic Wolbachia bacteria specifically inhibits oogenesis in a parasitic wasp ». Proceedings of the National Academy of Sciences 98 (11): 6247–6252.

Dupas, S., C.W. Gitau, A. Branca, B.P. Le Rü, & J.F. Silvain. 2008. « Evolution of a polydnavirus gene in relation to parasitoid–host species immune resistance ». Journal of Heredity 99 (5): 491.

Dupas, S., B. Le Ru, A. Branca, N. Faure, G. Gigot, P. Campagne, M. Sezonlin, R. Ndemah, G. Ong’amo, P.-A. Calatayud & J.-F. Silvain. 2014. « Phylogeography in continuous space: coupling species distribution models and circuit theory to assess the effect of contiguous migration at different climatic periods on genetic differentiation in Busseola fusca (Lepidoptera: Noctuidae) ». Molecular ecology 23 (9): 2313–25.

Dupuy, C., E. Huguet, & J.-M. Drezen. 2006. « Unfolding the evolutionary story of polydnaviruses ». Virus Research 117 (1): 81–89.

Durand, E., F. Jay, O. E. Gaggiotti, & O. François. 2009. « Spatial inference of admixture proportions and secondary contact zones ». Molecular Biology and Evolution 26 (9): 1963–1973.

Evanno, G., S. Regnaut, & J. Goudet. 2005. « Detecting the number of clusters of individuals using the software STRUCTURE: a simulation study ». Molecular ecology 14 (8): 2611–2620.

Excoffier, L., P. E. Smouse, & J. M. Quattro. 1992. « Analysis of molecular variance inferred from metric distances among DNA haplotypes: application to human mitochondrial DNA restriction data ». Genetics 131 (2): 479–491.

Fraley, C., A. E. Raftery, & L. Scrucca. 2012. « Normal mixture modeling for model-based clustering, classification, and density estimation ». Department of Statistics, University of Washington 23:2012.

Fraley, C., & A. E. Raftery. 2002. « Model-based clustering, discriminant analysis, and density estimation ». Journal of the American statistical Association 97 (458): 611–631.

François, O., M. Currat, N. Ray, E. Han, L Excoffier, & J. Novembre. 2010. « Principal component analysis under population genetic models of range expansion and admixture ». Molecular biology and evolution 27 (6): 1257–1268.

Freilich, X, J D. Anadón, J. Bukala, O.a Calderon, R. Chakraborty, & S. Boissinot. 2016. « Comparative Phylogeography of Ethiopian anurans: impact of the Great Rift Valley and Pleistocene climate change ». BMC evolutionary biology 16 (1): 206.

Gao, H., S. Williamson, & C. D. Bustamante. 2007. « A Markov chain Monte Carlo approach for joint inference of population structure and inbreeding rates from multilocus genotype data ». Genetics 176 (3): 1635–1651.

Gitau, C. W., D. Gundersen-Rindal, M. Pedroni, P. J. Mbugi, & S. Dupas. 2007. « Differential expression of the CrV1 haemocyte inactivation-associated polydnavirus gene in the African maize stem borer Busseola fusca (Fuller) parasitized by two biotypes of the endoparasitoid Cotesia sesamiae (Cameron) ». Journal of Insect Physiology 53 (7): 676–84.

Gitau, C. W, F. Schulthess, & S. Dupas. 2010. « An association between host acceptance and virulence status of different populations of Cotesia sesamiae, a braconid larval parasitoid of lepidopteran cereal stemborers in Kenya ». Biological Control.

Grimaldi, D. 2005. Evolution of the Insects. Cambridge University Press.

Habel, J. C., L. Borghesio, W. D. Newmark, J. J. Day, L. Lens, M. Husemann, & W. Ulrich. 2015. « Evolution along the Great Rift Valley: phenotypic and genetic differentiation of East African white-eyes (Aves, Zosteropidae) ». Ecology and evolution 5 (21): 4849–4862.

Harvey, C. P. 2011. Understanding and managing diversity: international edition. Pearson Education (Us).

Heino, J., A. S. Melo, L. M. Bini, F. Altermatt, S. A. Al-Shami, D. G. Angeler, N. Bonada, et al. 2015. « A comparative analysis reveals weak relationships between ecological factors and beta diversity of stream insect metacommunities at two spatial levels ». Ecology and evolution 5 (6): 1235–1248.

Henry, L. M., B. D. Roitberg, & D. R. Gillespie. 2008. « Host-range evolution in Aphidius parasitoids: fidelity, virulence and fitness trade-offs on an ancestral host. » Evolution 62 (3): 689.

Hilgenboecker, K., P. Hammerstein, P. Schlattmann, A. Telschow, & J. H. Werren. 2008. « How many species are infected with Wolbachia-–a statistical analysis of current data ». FEMS Microbiology Letters 281 (2): 215–220.

Holt, R D, & M E Hochberg. 1997. « When is biological control stable (or is it)? » Ecology 78 (6): 1673–83.

Hoskin, C. J., & M. Higgie. 2010. « Speciation via species interactions: the divergence of mating traits within species ». Ecology letters 13 (4): 409–420.

Jaenike, J., K. A. Dyer, C. Cornish, & M. S. Minhas. 2006. « Asymmetrical reinforcement and Wolbachia infection in Drosophila ». PLoS Biol 4 (10): e325.

Jabot, F. & J. Bascompte. 2012. « Bitrophic interactions shape biodiversity in space ». Proceedings of the National Academy of Sciences 109 (12): 4521–4526.

Jakobsson, M., & N. A. Rosenberg. 2007. « CLUMPP: a cluster matching and permutation program for dealing with label switching and multimodality in analysis of population structure ». Bioinformatics 23 (14): 1801–1806.

Jancek, S., A. Bézier, P. Gayral, C. Paillusson, L. Kaiser, S. Dupas, B. P. Le Ru, et al. 2013. « Adaptive selection on bracovirus genomes drives the specialization of Cotesia parasitoid wasps. » PloS one 8 (5): e64432.

Jensen, M. K., K. M. Kester, M. Kankare, & B. L. Brown. 2002. « Characterization of microsatellite loci in the parasitoid, Cotesia congregata (Say)(Hymenoptera Braconidae) ». Molecular Ecology Notes 2 (3): 346–348.

Jombart, T., & I. Ahmed. 2011. « adegenet 1.3-1: new tools for the analysis of genome-wide SNP data ». Bioinformatics 27 (21): 3070–3071.

Kaiser, L., B. Le Ru, F. Kaoula, C. Paillusson, C. Capdevielle-Dulac, J. Obonyo, E. Herniou, S. Jancek, A. Branca, P.-A. Calatayud, J.-F. Silvain & S. Dupas. 2015. « Ongoing ecological speciation in Cotesia sesamiae, a biological control agent of cereal stemborers ». Evolutionary Applications 8(8), 807–820.

Kaiser, L., J. Fernandez-Triana, C. Capdevielle-Dulac, C. Chantre, M. Bodet, F. Kaoula, R. Benoist, et al. 2017. « Systematics and biology of Cotesia typhae sp. n.(Hymenoptera, Braconidae, Microgastrinae), a potential biological control agent against the noctuid Mediterranean corn borer, Sesamia nonagrioides ». ZooKeys 682: 105.

Kfir, R. 1995. « Parasitoids of the African stem borer, Busseola fusca (Lepidoptera: Noctuidae), in South Africa ». Bulletin of Entomological Research 85 (03): 369–77.

Kfir, R., W. A. Overholt, Z. R. Khan, & A. Polaszek. 2002. « Biology and management of economically important lepidopteran cereal stem borers in Africa. » Annual review of entomology 47: 701–31.

Kimani-Njogu, S. W., W. A. Overholt, J. Woolley, & A. Walker. 1997. « Biosystematics of the Cotesia flavipes species complex (Hymenoptera: Braconidae): morphometrics of selected allopatric populations ». Bulletin of Entomological Research 87 (1): 61–66.

Kottek, M., J. Grieser, C. Beck, B. Rudolf, & F. Rubel. 2006. « World map of the Koppen-Geiger climate classification updated ». Meteorologische Zeitschrift 15 (3): 259–264.

Le Ru, B. P., G. O. Ong’amo, P. Moyal, E. Muchugu, L. Ngala, B. Musyoka, Z. Abdullah, et al. 2006. « Geographic distribution and host plant ranges of East African noctuid stem borers ». In Annales de la Société entomologique de France, 42:353–361.

Mairal, M., I. Sanmartín, A. Herrero, L. Pokorny, P. Vargas, J. J. Aldasoro, & M. Alarcón. 2017. « Geographic barriers and Pleistocene climate change shaped patterns of genetic variation in the Eastern Afromontane biodiversity hotspot ». Scientific Reports 7.

McArdle, B. H., & M. J. Anderson. 2001. « Fitting multivariate models to community data: a comment on distance-based redundancy analysis ». Ecology 82 (1): 290–297.

Meirmans, P. G., & P. H. Van Tienderen. 2004. « GENOTYPE and GENODIVE: two programs for the analysis of genetic diversity of asexual organisms ». Molecular Ecology Notes 4 (4): 792–794.

Mellersh, C., & J. Sampson. 1993. « Simplifying detection of microsatellite length polymorphisms ». BioTechniques 15 (4): 582–584.

Mochiah, M. B., A. J. Ngi-Song, W. A. Overholt, & R. Stouthamer. 2002a. « Variation in encapsulation sensitivity of Cotesia sesamiae biotypes to Busseola fusca ». Entomologia Experimentalis et Applicata 105 (2-3): 11–118.

Mochiah, M. B, A. J. Ngi-Song, W. A. Overholt, & R. Stouthamer. 2002b. « Wolbachia infection in Cotesia sesamiae (Hymenoptera: Braconidae) causes cytoplasmic incompatibility: implications for biological control ». Biological Control 25 (1): 74–80.

Ngi-Song, A. J., W. A. Overholt, & J. N. Ayertey. 1995. « Suitability of African gramineous stemborers for development of Cotesia flavipes and C. sesamiae (Hymenoptera: Braconidae) ». Environmental Entomology 24.

Ngi-Song, A. J., & M. B. Mochiah. 2001. « Polymorphism for Wolbachia infections in eastern and southern African Cotesia sesamiae (Cameron)(Hymenoptera: Braconidae) populations ». International Journal of Tropical Insect Science 21 (04): 369–374.

Nuismer, S. L. 2006. « Parasite local adaptation in a geographic mosaic. » Evolution; international journal of organic evolution 60 (1): 24–30.

Ode, P., M. Antolin, & M. Strand. 1998. « Differential dispersal and female-biased sex allocation in a parasitic wasp ». Ecological Entomology 23 (3): 314–318.

Odee, D. W., A. Telford, J. Wilson, A. Gaye, & S. Cavers. 2012. « Plio-Pleistocene history and phylogeography of Acacia senegal in dry woodlands and savannahs of sub-Saharan tropical Africa: evidence of early colonisation and recent range expansion ». Heredity 109 (6): 372.

Oksanen, J., F. G. Blanchet, R. Kindt, P. Legendre, P. R. Minchin, R. B. O’Hara, G. L. Simpson, M. J. Oksanen, & M. Suggests. 2013. « Package ‘vegan’ ». http://mirror.bjtu.edu.cn/cran/web/packages/vegan/vegan.pdf.

Ong’amo, G. O., B. P. Le Ru, S. Dupas, P. Moyal, P.-A. Calatayud, & J.-F. Silvain. 2006. « Distribution, pest status and agro-climatic preferences of lepidopteran stem borers of maize in Kenya ». Annales de la Société Entomologique de France 42: 171–178.

Onyango, F. O., & J. P. R. Ochieng’-Odero. 1994. « Continuous rearing of the maize stem borer Busseola fusca on an artificial diet ». Entomologia Experimentalis et Applicata 73 (2): 139–144.

Overholt, W. A., J. O. Ochieng, P. Lammers, & K. Ogedah. 1994. « Rearing and field release methods for Cotesia flavipes Cameron (Hymenoptera: Braconidae), a parasitoid of tropical gramineous stem borers ». International Journal of Tropical Insect Science 15 (03): 253–259.

Potting, R. P. J., N. E. Vermeulen, & D. E. Conlong. 1999. « Active defence of herbivorous hosts against parasitism: Adult parasitoid mortality risk involved in attacking a concealed stemboring host ». In Proceedings of the 10th International Symposium on Insect-Plant Relationships, 143–148. Springer.

Roderick, G. K.. 1996. « Geographic structure of insect populations: gene flow, phylogeography, and their uses ». Annual review of entomology 41 (1): 325–352.

Santos, A. M. C., & D. L. J. Quicke. 2011. « Large-Scale Diversity Patterns of Parasitoid Insects ». Entomological Science 14 (4): 371–382.

Sepulchre, P., G. Ramstein, F. Fluteau, M. Schuster, J.-J. Tiercelin, & M. Brunet. 2006. « Tectonic uplift and Eastern Africa aridification ». Science 313 (5792): 1419–1423.

Sezonlin, Michel, Stéphane Dupas, B. Le Rü, Philippe Le Gall, Pascal Moyal, P.-A. Calatayud, I. Giffard, N. Faure, & J.-F. Silvain. 2006. « Phylogeography and population genetics of the maize stalk borer Busseola fusca (Lepidoptera, Noctuidae) in sub-Saharan Africa ». Molecular Ecology 15 (2): 407–420.

Smouse, P. E., & R. Peakall. 1999. « Spatial autocorrelation analysis of individual multiallele and multilocus genetic structure ». Heredity 82 (5): 561–573.

Stiling, P., & T. Cornelissen. 2005. « What makes a successful biocontrol agent-A meta-analysis of biological control agent performance ». Biological Control 34 (3): 236–46.

Telschow, A., J. Engelstädter, N. Yamamura, P. Hammerstein, & G. D. D. Hurst. 2006. « Asymmetric gene flow and constraints on adaptation caused by sex ratio distorters ». Journal of Evolutionary Biology 19 (3): 869–878.

Thompson, J N.. 2005. The geographic mosaic of coevolution. Chicago: The university of Chicago Press.

Ullyett, G. C.. 1935. « Notes on Apanteles sesamiae Cam., a parasite of the maize stalk-borer (Busseola fusca, Fuller) in South Africa ». Bulletin of Entomological Research 26 (02): 253–262.

Van Nouhuys, S., & I. Hanski. 2002. « Colonization rates and distances of a host butterfly and two specific parasitoids in a fragmented landscape ». Journal of Animal Ecology 71 (4): 639–650.

Van Valen, L.. 1973. « A new evolutionary law ». Evolutionary Theory 1 (1): 1–30.

Vavre, F., F. Fleury, J. Varaldi, P. Fouillet, & M. Boulétreau. 2000. « Evidence for female mortality in Wolbachia-mediated cytoplasmic incompatibility in haplodiploid insects: epidemiologic and evolutionary consequences ». Evolution, 191–200.

Werren, J H.. 1997. « Biology of Wolbachia. » Annual review of entomology 42 (124): 587–609.

Whitfield, J. B.. 2002. « Estimating the age of the polydnavirus/braconid wasp symbiosis ». Proceedings of the National Academy of Sciences of the United States of America 99 (11): 7508–13.

Zhou, Y., H. Gu, & S. Dorn. 2006. « Single-locus sex determination in the parasitoid wasp Cotesia glomerata (Hymenoptera: Braconidae) ». Heredity 96 (6): 487–492.

